# Antibiotics increase aggression behavior and aggression-related pheromones and receptors in *Drosophila melanogaster*

**DOI:** 10.1101/2020.09.22.307777

**Authors:** Michal Grinberg, Hadar Neuman, Oren Ziv, Sondra Turjeman, Rita Nosenko, Omry Koren

## Abstract

Aggression is a behavior common in most species; it is controlled by internal and external drivers, including hormones, environmental cues, and social interactions, and underlying pathways are understood in a broad range of species. To date, though, effects of gut microbiota on aggression in the context of gut-brain communication and social behavior have not been elucidated. We examine how manipulation of *Drosophila melanogaster* microbiota affect aggression as well as the pathways that underly the behavior in this species. Flies treated with antibiotics exhibited significantly more aggressive behaviors. Furthermore, they had higher levels of vCA and (Z)-9 Tricosene, pheromones associated with aggression in flies, as well as higher expression of the relevant pheromone receptors and transporters OR67d, OR83b, GR32a, and LUSH. These findings suggest that aggressive behavior is, at least in part, mediated by bacterial species in flies.

## Introduction

Aggression is evident in almost all animal species and can be influenced by specific genes, neurotransmitters, neural systems, pheromones, hormones, social interactions, and other environmental factors^1, 2^. Aggressive behavior and pathways controlling it are well studied in model organisms ^3-7^, but a nuanced understanding of how certain biological processes interact with these pathways is lacking. Specifically, it is evident that the gut microbiota can greatly influence aspects of host physiology, including gut–brain communication and social behavior^8, 9^, but to date, the effect of the gut microbiota on aggression and underlying pathways is not fully understood. In the current study, we asked whether the microbiome plays a role in aggressive behavior and if so, what pathways may be involved.

Here we focus on *Drosophila melanogaster* because aggression has been well-studied in this simple animal model^10^. The neuronal mechanisms leading to aggression in *D. melanogaster* have been identified and mainly include pheromones and olfactory sensory neurons that express odorant receptors^11^. Furthermore, previous studies of gut-brain-behavior interactions in this species demonstrated a clear influence of antibiotics on mating preference, correlated with alterations in cVA levels^8^; cVA is a male-specific pheromone known to affect courtship and aggression in fruit flies^12^. Additionally, the gut endosymbiont bacteria, *Wolbachia*, was shown to alter pheromone production in *D. melanogaster* pupae, interfering with their communication and causing gamete incompatibility^13^. Due to its relatively simple microbiota composition and the ability to easily manipulate it to test the effects of single bacterial species on behavior, *D. melanogaster* is currently one of the preferred model animals in the field of gut–brain communication and social behavior^8, 14-16^.

Previous studies have confirmed that the microbiota plays a role in gut–brain communication and subsequent behavior^8, 9^. Accordingly, we hypothesized that alterations in the fly microbiome would affect male aggression behavior by modulating expression of the related pheromones (cVA, (Z)-9 Tricosene)^11, 17, 18^ and receptor and transporter components (OR67d, OR83b, GR32a, LUSH)^19-21^. Through a set of manipulations, we studied how specific changes to microbiota composition alter aggression behavior and examined how the microbiota interacts with relevant pheromones and receptors.

## Results

To test our overarching hypothesis that microbiome alterations affect male fly aggression, we measured aggression^2, 13^ in four experimental groups of male *D. melanogaster*: (1) **untreated** flies (control group), (2) **Abx** flies grown on media supplemented with a mixture of antibiotics (to eliminate gut bacteria; Supp. Fig. 1), (3) ***Lactobacillus brevis***-monocolonized flies, and (4) ***L. plantarum***-monocolonized flies. The flies in groups 3 and 4 were offspring of flies grown on antibiotics that were transferred to sterile media supplemented the focal microbe. We found that Abx treatment increased the number of aggressive encounters among male flies compared to the control group by nearly 150% (Fig. 1; *p*-value < 0.05) whereas supplementation with a single bacterial species (*L. plantarum* or *L. brevis*) reduced aggressive behavior compared to both the Abx-treated flies (Fig 1; *p*-value < 0.0001 and < 0.005 respectively) and the control group (marginally significant, *p*-value = 0.09). These results validated our hypothesis that bacteria can modulate aggression behavior.

**Figure 1.**
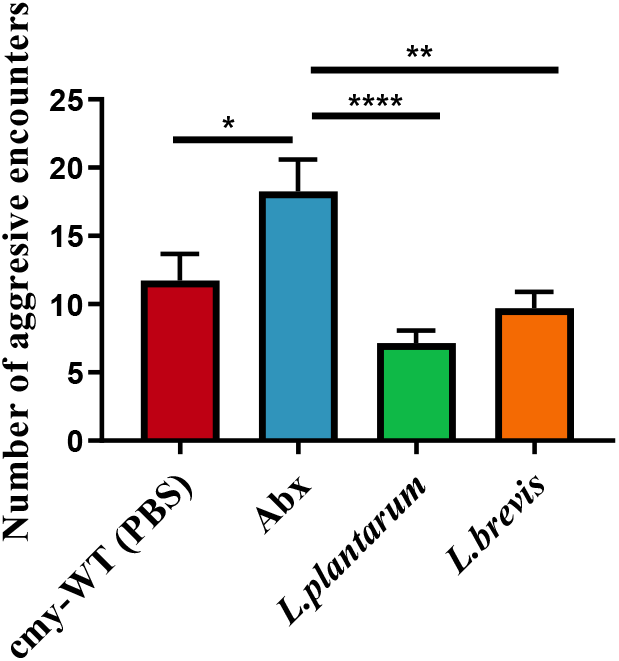
Aggression levels are influenced by microbial changes in *D. melanogaster*. Behavior tests showing the different number of aggressive encounters in the four treatment groups. The Abx treated flies showed higher levels of aggression than any other treatment, while treatment with a monoculture of *L. plantarum* proved to reduce the aggression levels the most (n=20 repetitions per treatment). Statistics calculated with one-way ANOVA; *< 0.05, **<0.005, ****<0.0001; bars indicate S.E.

To decipher the mechanism underlying this interaction, we first examined how the gut microbiota influences levels of cVA and (Z)-9 Tricosene (9-T), pheromones typically positively associated with aggressive behavior in male fruit flies^22^. Using the same experimental set up, we found that levels of both pheromones were significantly higher in Abx-treated flies than in other treatment groups (Fig. 2a,b); specifically, cVA and 9-T levels were on average 2 times greater than the control, respectively.

**Figure 2.**
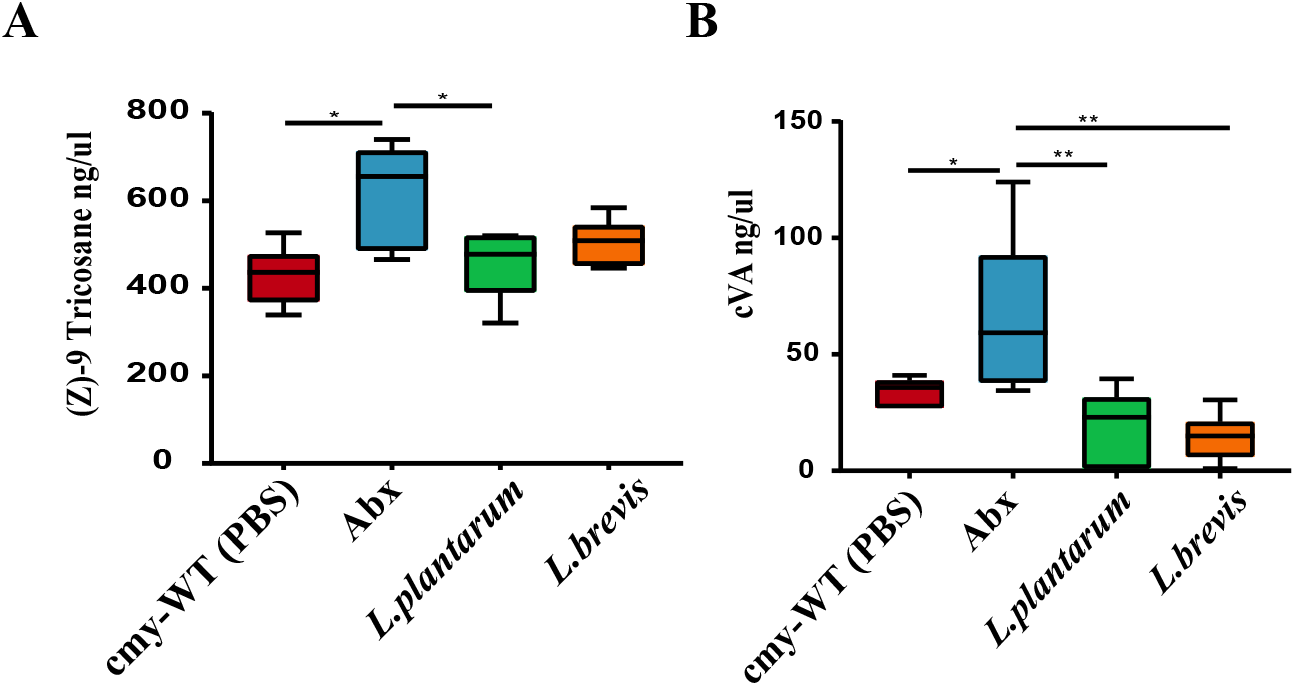
Levels of pheromone changes according to the microbiome composition. (A) (B) Levels of aggressive pheromones in male flies. Pheromone levels calculated by concentration of internal standard with GC-MS (n=8 vials with 8 male flies for each treatment). (A) Expression of (Z)-9 Tricosane levels by treatment. Higher levels measured in Abx treated flies and lower levels in *L. plantarum* monocultures. (B) Expression of cVA levels by treatment. Higher levels measured in Abx treated flies and lower levels in *L. plantarum* and *L. brevis* monocultures. Statistics calculated with one-way ANOVA; *< 0.05, **<0.005; bars indicate S.E.

We next examined the effect of microbiota on three pheromone receptors associated with aggression, OR67d and OR83b, receptors of cVA^17^, and GR32a, a receptor of 9-T^23^, using qRT-PCR to quantify their expression (Fig. 3a-c). Abx treatment significantly raised levels of OR83b and GR32a, but not OR67d, compared to untreated flies. Interestingly, *L. plantarum* supplementation significantly and most dramatically raised levels of all three receptors whereas *L. brevis* supplementation resulted in receptor levels comparable to or slightly greater than untreated flies. In addition to measurements of receptor expression, we also compared levels of the cVA transporter LUSH between groups^21^ (Fig. 3d). Again, the Abx treatment resulted in significantly higher expression levels compared to all other groups.

**Figure 3.**
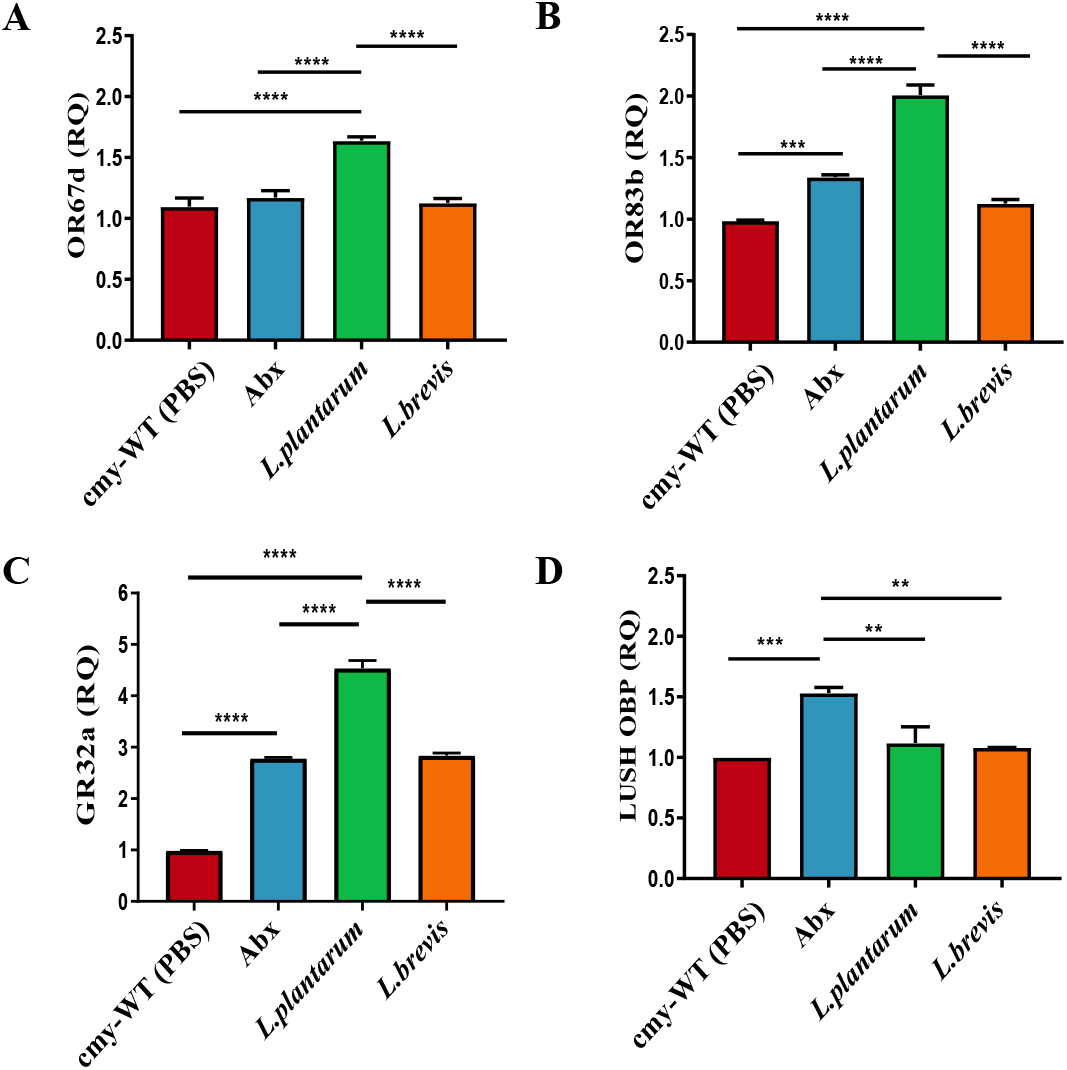
RNA expression of OSN in the different groups. Levels of RNA were calculated using qPCR and obtaining the RQ value in each group (n=30 male flies; 3 biological repetitions). **A)** OR67d. **B)** OR83b (ORCO). **C)**. In *L. plantarum* monocultured flies, the expression was higher than other treatments in GR32a. **D)** Higher levels were observed in Abx treatment in LUSH OBP expression. Statistics calculated by one-way ANOVA; **< 0.005, ***<0.001, ****<0.0001; bars indicate S.E.

## Discussion

We tested the hypothesis that fruit fly microbiota affects male aggression behavior, with behavioral tests as well as experiments that examined pheromone production and receptor/transporter expression. This hierarchical study design allowed us to unravel the cascade of effects the microbiota has on the physiology of aggression by approaching the pathway holistically. Combined, our findings show that Abx treatment increased aggression in male flies, as compared to untreated flies, with Abx-flies producing higher levels of aggression pheromones cVA and (Z)-9 Tricosene and exhibiting higher expression of the related receptor components OR83b and GR32a and cVA transporter LUSH. We thus conclude that the natural gut microbiota in the fly plays a role in regulating male aggression by both modulating pheromone production as well as expression of their receptors and associated proteins.

Of note, while Abx treatment significantly raised levels of OR83b and GR32a, as compared to untreated flies, it did not seem to influence OR67d, suggesting that this may be a less important receptor of vCA than OR83b. Also of interest, in the treatment group supplemented with *L. plantarum*, we found low levels of aggression accompanied by low pheromone levels, yet receptor levels were significantly higher than in all other groups. This phenomenon might be explained by a negative feedback loop previously described in this pathway^20, 24^.

Our study is the first to show the relationship between antibiotics, aggression, and also pheromones and receptor levels. Understanding these relationships can provide more information about gut-brain communication necessary for deciphering behavioral mechanisms related to aggression as well as additional behaviors. Further monocolonization studies can uncover the nuances associated with this behavior, and bacterial species found to moderate aggression can be examined for similar interactions in other species.

## Materials and Methods

### Fly stocks

Fly stocks (Oregon R) were obtained from Bloomington Drosophila Stock Center (Indiana Avenue, Bloomington, IN, USA). Flies were reared in 50 ml vials containing 10 ml CMY (cornmeal, molasses, yeast) growth media. Flies were maintained in an incubator at 25°C with a light dark cycle of 12h:12h.

### Antibiotic treatment (Abx)

An antibiotic mixture containing three types of antibiotics (50 µg/ml tetracycline, 200 µg/ml rifampicin, and 100 µg/ml streptomycin) was added to CMY media. In order to functionally test the effectiveness of the Abx supplementation, a PCR using microbial primers for the 16S rRNA gene (515F+806R)^25^ was performed and showed absence of microbial DNA.

### Single microbe supplementation

Abx-treated flies were transferred to new vials containing CMY media supplemented with 100 µl of an overnight culture (∼10^8^ bacteria) of either *L. plantarum* or *L. brevis* diluted in sterile PBS. The bacterial concentration chosen is comparable to levels in our untreated flies. Offspring of transferred flies (2^nd^ generation) were used for experiments.

### Aggression experiment

Eight male flies aged 4-7 days were collected in empty vials and starved for 2 hours^22^. The flies were then transferred (without anesthesia) to a vial containing a patch of yeast-water and a decapitated female, which provide ideal conditions for aggression^22^. For the first 5 min, the flies were left to adapt to the new vial. Their activity was then recorded for 5 min using a video camera. The total number of aggression encounters within the vial (including chasing, wing threat, boxing and lunging) was recorded. At least 20 aggression tests were analyzed per experimental group. Scoring was performed in a blinded manner.

### Gas chromatography analysis

Eight flies aged 4-7 days were separated into an empty glass vial and starved for 2 hours. Pheromones were then extracted from fly cuticles by adding 1000 µl hexane for 5 min at room temperature. The liquid was transferred to a GC-MS adjusted vial, and 10 µl of hexane containing 2000 ng/µl of hexocosane (C-26) was added as an internal standard. Vials were shaken for 1 min. Extracts were concentrated to 10 µl, of which 2 µl were injected into a HP-5/mS silica capillary column (30m*0.25mm*0.25mm film thickness) that was temperature-programmed: 140°C (2 min), 3°Cmin^−1^ to 300°C (2 min). Extracts were analyzed by gas chromatography coupled with mass spectrometry (Clarus SQ 8T GC/ Mass Spectrometer, Perkin Elmer, Walthman, MA, USA). The NIST mass-spectral library identifications were confirmed with chemical standards when available (Sigma-Aldrich). Compounds of interest (cVA and (Z)-9 Tricosene) were identified based on their mass spectrum and retention time and quantified by peak integration. These two compounds were chosen because they have previously been identified as pheromones associated with aggression in male flies^22^.

### qRT-PCR

For each treatment group, ten male flies (4-7 days old) were collected and anesthetized. Decapitation was performed using sterile tweezers. RNA was purified following homogenization of the heads with a total RNA purification kit according to the manufacturer’s protocol (NORGEN, Thorold, ON, Canada). The first strand of cDNA (see protein targets below) was synthesized from 5X single RT MasterMix (abm, Vancouver, BC, Canada) using reverse transcriptase. Quantitative real-time PCR was performed using the StepOne™ Real-Time PCR System (Thermo Fisher Scientific, Waltham, MA, USA). Reactions included a mixture of 5 µl 2X SYBR, 1 µl of each primer (10 µM; see below), and 4 µl cDNA per sample. In the negative control, cDNA was replaced with DDW. Primers were designed using Primer-BLAST (NCBI) and FlyPrimerBank (DRSC; https://www.flyrnai.org/FlyPrimerBank) for well-studied pheromone receptors and a transporter as targets: OR67d (receptor for vCA), OR83b (vcA), GR32a (9-T), and LUSH (transporter for vCA)^18^.

OR67d F: ATTTTGCGGAAACGATGTGGC R: GGATTATGGTGAGGTCTCCATTG

OR83b F: TCACGAAGTTTATCTACCTGGCT R: ATCGAATGGTAACGAGCATCC

GR32a F: CTATGAGGTGGGTCCTCCGA R: CGTCTCGCGGTAGGAGAAAA

LUSH F: GACCTCGCTAGACATGATCCG R: GCACATAAGATCCTGCGATGG

### Data analysis

Two-tailed one-way ANOVAs were used to test differences in aggressive encounters and pheromone and related protein expression levels among the four experimental treatments.

## Funding

This study was funded by the Israel Science Foundation, Grant Numbers 2459/15 and 1001/16.

## Notes

### Competing Interest Statement

The authors have declared no competing interest.

